# Neonatal granulocytic MDSCs possess phagocytic properties during bacterial infection

**DOI:** 10.1101/2019.12.30.891077

**Authors:** Brittany G. Seman, Jordan K. Vance, Michelle R. Witt, Cory M. Robinson

**Affiliations:** Department of Microbiology, Immunology, & Cell Biology, West Virginia University School of Medicine, Morgantown, WV; Vaccine Development Center at West Virginia University Health Sciences Center, Morgantown, WV, USA

## Abstract

Myeloid-derived suppressor cells (MDSCs) are an immunosuppressive cell type found in high abundance in early life. Currently, there has been limited mechanistic understanding of MDSC phagocytosis of bacteria and the corresponding consequences in the context of acute infection. We set out to determine whether human granulocytic MDSCs have phagocytic capacity that is comparable to other professional phagocytes. To investigate these properties, we utilized fluorescent confocal microscopy, flow cytometry, and bacterial burden assays. We demonstrate that human granulocytic MDSCs phagocytose *E. coli* O1:K1:H7, and subsequently traffic the bacteria into acidic compartments similar to other phagocytes. However, MDSCs were significantly less efficient at bacterial uptake and killing compared to monocytes. This activity is associated with an inflammatory response, but the amount of TNFα gene and protein expression was reduced in infected MDSCs compared to monocytes. Interestingly, we also found that MDSCs release DNA (MeDNA) into the extracellular space that resembles neutrophil extracellular traps. We found that MeDNA had some impact on bacterial viability in single cultures, with an increase in bacterial recovery in MDSCs treated with DNAse. However, MeDNA did not impact the ability of monocytes to eliminate bacteria in co-cultures, suggesting that MDSC extracellular DNA does not compromise monocyte function. Overall, our data reveals mechanistic insight into MDSC activity during infection that includes the kinetics and efficiency of bacterial uptake, elimination through trafficking to acidified compartments, and inflammatory contributions relative to primary human monocytes. These results enhance our understanding of MDSC contributions during acute bacterial infection and identify host-directed targets for immune intervention to improve outcomes and reduce susceptibility to infection early in life.

## Introduction

Myeloid-derived suppressor cells are an immune suppressive cell type found in high abundance during the unique immune period of early life (Gervassi et al., 2014; Rieber et al., 2013). Originally, MDSCs were observed to promote cancer progression by suppressing anti-tumor immunity and compromising T cell surveillance (Bronte et al., 2000; Young, Newby, & Wepsic, 1987). Interestingly, the abundance of MDSCs during early life correlates with an increased susceptibility of neonates to infection and mortality due to infection (Schwarz et al., 2018). However, there is limited data on MDSC interactions with bacteria and the innate immune control of acute infection in the context of early life immunity. Thus, it is necessary to evaluate the direct interactions of MDSCs with bacteria and subsequent immunoregulatory activity. This may lead to the identification of novel host-targeted therapies for early life infections.

The neonatal immune system is characterized by a distinct, regulatory state compared to adults. Although there are increases in T cells, B cells, neutrophils, and monocytes in neonates compared to adults (Sharma, Jen, Butler, & Lavoie, 2012), there are differences in activity and function relative to adult counterparts (Simon, Hollander, & McMichael, 2015). For instance, neonatal neutrophils are defective in L-selectin and CD11b production, which impairs migration to sites of infection and wounding (Kim, 2003; O’Hare et al., 2015). Neonatal monocytes have lower levels of costimulatory and antigen-presenting molecules such as HLA-DR, CD86, and CD40 compared to adults, which aids in the delayed response to stimulatory molecules such as lipopolysaccharide (LPS) (Velilla, Rugeles, & Chougnet, 2006). The neonatal immune system is also polarized to a more regulatory Th2 state (Marodi, 2002; Protonotariou et al., 2010). Taken together, these reported findings suggest that the neonatal immune system exhibits an altered profile compared to adults that is less equipped to combat bacterial infections.

Another unique feature of the neonatal immune system is the abundance of MDSCs. Circulating MDSCs are found at higher numbers in umbilical cord blood than peripheral blood of children and adults (Rieber et al., 2013; Schwarz et al., 2018). MDSCs are also approximately two-fold more abundant in neonatal mouse spleens compared with adults (Gleave Parson et al., 2019). MDSCs have important immunosuppressive functions during disease. During tumor growth, the expression of CD40 on MDSCs has been shown to induce tumor tolerance, as well as increase Treg production (Pan et al., 2010). MDSCs also suppress NK cell production of interferon-gamma (IFNγ) through cell-to-cell contact and tumor growth factor-beta (TGF-β) production, subsequently affecting NK antitumor immunity (H. Li, Han, Guo, Zhang, & Cao, 2009). MDSCs are separated into two main subsets: granulocytic (gMDSCs) and monocytic (mMDSCs) (Peranzoni et al., 2010). Classical effector functions linked with immune suppression include the production of arginase, nitric oxide, and reactive oxygen species. Collectively, these factors suppress multiple immune cell types through depletion of arginine, the inhibition of JAK-STAT proteins, and the reduction of MHC class II expression (Darcy et al., 2014; Gabrilovich & Nagaraj, 2009). MDSCs express a multitude of both proinflammatory and anti-inflammatory cytokines, including tumor necrosis factor alpha (TNFα) and interleukin-10 (IL-10) (Janols et al., 2014; Poe et al., 2013). Our lab has also recently determined MDSCs to be a source of IL-27, a cytokine known for its suppression of inflammation (Gleave Parson et al., 2019). Taken together, this body of literature suggests that MDSCs are not only abundant in the neonatal immune system, but are also important suppressive immune cell types during disease in both adults and neonates.

MDSCs have been directly implicated in the altered function of other immune cells during infection. Our lab has demonstrated that macrophages are impaired in their ability to clear bacteria *in vitro* in the presence of MDSCs (Gleave Parson et al., 2019). Monocytes from human umbilical cord blood are impaired in their ability to stimulate T cell activation and phagocytose bacteria due to the influence of MDSCs (Dietz et al., 2019). MDSCs have also been involved in the immune shift towards an anti-inflammatory state during late-onset sepsis induced by cecal ligation and puncture in mice (Brudecki, Ferguson, McCall, & El Gazzar, 2012). However, few studies have investigated the direct interactions that occur between MDSCs and microbes during acute infection. Although MDSCs share a common progenitor with professional phagocytes, the phagocytic capabilities of MDSCs have yet to be fully analyzed. Davis and colleagues briefly suggested that MDSCs can phagocytose *Escherichia coli* (*E. coli*) particles *in vitro*; however, the mechanistic details were not analyzed in depth (Davis, Silvin, & Allen, 2017). Additionally, two other studies reported that MDSCs can internalize *Mycobacterium tuberculosis* and *Mycobacterium bovis* Bacillus Calmette-Guerin (BCG), although the mechanisms, kinetics, and efficiency of internalization were not addressed (Agrawal et al., 2018; Magcwebeba, Dorhoi, & du Plessis, 2019; Martino et al., 2010). As such, our mechanistic understanding of direct MDSC interactions with bacterial pathogens has remained limited, and the consequences during infection have been unclear.

Herein we describe the phagocytic capabilities of human granulocytic MDSCs during bacterial infection *in vitro*. Using the pathogenic *E. coli* serotype O1:K1:H7, which is a leading cause of invasive neonatal infections such as sepsis and meningitis that are responsible for significant mortality (Simonsen, Anderson-Berry, Delair, & Davies, 2014; Stoll et al., 2011), we examined the phagocytic capacity of MDSCs through 3D time lapse microscopy, flow cytometry, and bacterial killing assays. Our data suggests that MDSCs are capable of phagocytic uptake and elimination of bacteria, although they are functionally limited compared to monocytes. Additionally, we find that MDSCs release DNA into the extracellular environment, although this function is not associated with potent bactericidal activity. In contrast, bacterial clearance by monocytes co-cultured with MDSCs is improved in the absence of extracellular DNA. This study demonstrates novel MDSC functionality that has not been rigorously evaluated, and gives rise to new questions surrounding the contributions of MDSCs in the host response during acute infections.

## Results

### MDSCs have the ability to phagocytose bacteria

MDSCs are well characterized by their ability to suppress immunity, and are functionally defined by their ability to limit T cell proliferation (Chen et al., 2017; Ostrand-Rosenberg, Sinha, Beury, & Clements, 2012). However, much less is known about how they interact directly with bacterial pathogens or within the overall immune response during infection. To test whether MDSCs have the capacity to phagocytose during infection, we isolated granulocytic MDSCs from human umbilical cord blood. Granulocytic MDSCs were separated from mature neutrophils by density gradient separation. This isolation strategy yielded CD66^hi^, CD33^+^, CD14^lo^ and HLA-DR^-^ granulocytic cells that suppress IL-2- and CD3/CD28 DynaBead-induced CD4^+^ T cell proliferation (Supplemental Figure 1). *E. coli* O1:K1:H7 was labeled with Syto 9 and cultured with MDSCs at increasing multiplicities of infection (MOI) for 1.5-2 hours. The MDSCs were then examined by flow cytometry for fluorescent cells. Fluorescent events increased dose-dependently with increasing MOI, suggesting that MDSCs do have the capacity to internalize bacteria (Figure 1A). Indeed, phagocytosis of bacteria directly correlated with increasing MOI (Figure 1B). To further examine the phagocytosis of bacteria while excluding any extracellular-associated bacteria, *E. coli* were conjugated with the pH-sensitive dye, pHrodo Red. Using time lapse microscopy, we found that MDSCs contain acidic compartments in which internalized bacteria were localized to, with internalization occurring as quickly as 8 minutes into imaging (Figure 1C). Taken together, these results demonstrate that granulocytic MDSCs have the ability to phagocytose bacteria and shuttle them into acidic compartments, similar to professional phagocytes.

**Figure 1:**
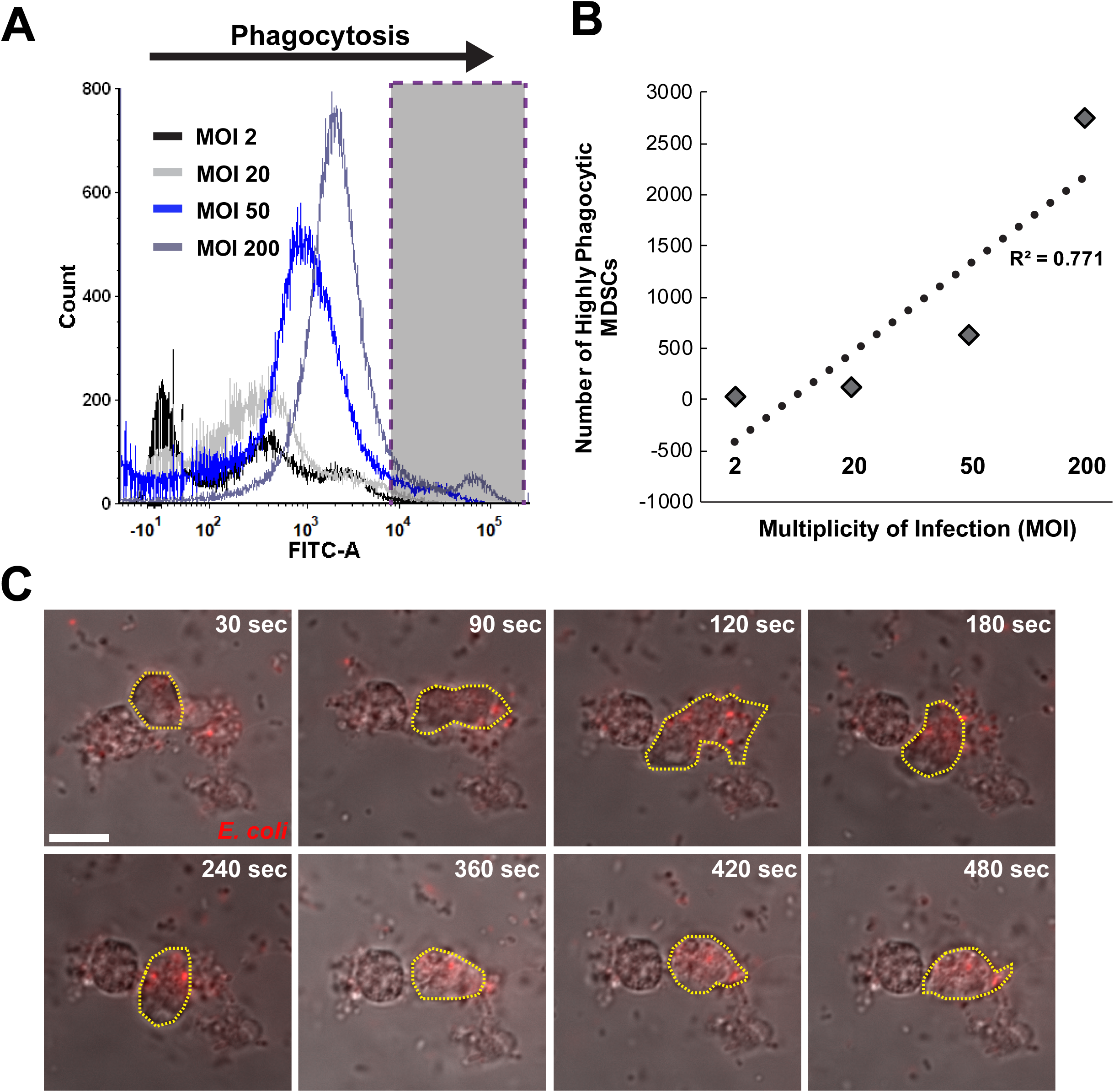
MDSCs phagocytose bacteria in a dose-dependent manner. CD66^+^ MDSCs isolated from human umbilical cord blood PBMCs were infected with varying multiplicities of infection (MOI) of pHrodo™ Red- or Syto 9™-labeled *E. coli* O1:K1:H7 and incubated at 37°C. For imaging, cells were longitudinally imaged over 10-20 minutes on a Nikon A1R confocal microscope at 40X to capture phagocytosis in real time. For flow cytometry, cells were fixed in 4% paraformaldehyde and resuspended in PBS prior to collection. **(A, B, and C)** Images and graphs are representative of 3 independent experiments. **(A)** A representative infection histogram overlay of four bacterial MOIs is shown. Bacteria labeled with Syto 9™ were quantified from gated MDSCs. MOI 2 = black line, MOI 20 = light grey line, MOI 50 = royal blue line, MOI 200 = blue-grey line. Grey box with purple dotted outline is the area of highly phagocytic MDSCs. **(B)** A scatter plot with best fit line correlating MOI to the number of highly phagocytic MDSCs is shown. Each dark grey diamond symbol represents an average number of phagocytic MDSCs per MOI. The R^2^ value is displayed. **(C)** Representative images of MDSC phagocytosis of pHrodo™ Red-labeled bacteria over a period of 15 minutes are shown. An MDSC of interest is outlined with a yellow dotted line. Panels are captured at time points described in seconds in the top right of images. Scale bar = 100 μm.

### MDSCs are less efficient at uptake and elimination of bacteria than monocytes

MDSCs are able to phagocytose and internalize bacteria into acidic compartments. Next, we wanted to determine the kinetics and efficiency of this activity relative to professional phagocytes. Neutrophils have limited longevity, and since we are unable to receive umbilical cord blood immediately post-collection, reduced viability between collection and transit to the lab may affect the interpretation of later time points during experiments. Therefore, for these experiments, we isolated and used human CD14^+^ monocytes, which are readily available in the same peripheral blood mononuclear cell (PBMC) fraction. To determine the ability of MDSCs to eliminate bacteria upon phagocytosis, we infected cells with fluorescently-labeled *E. coli* and longitudinally quantified uptake and bacterial recovery compared to monocytes. Overall, we found that MDSCs are significantly less efficient at bacterial uptake compared to monocytes. Figure 2A illustrates a significant increase in the uptake of large bacterial quantities by monocytes compared to MDSCs during flow cytometry. This increase in bacterial uptake was also quantified by confocal microscopy as the number of fluorescent pHrodo particles phagocytosed per cell (Figure 2B), as well as the area of pHrodo fluorescence per image between MDSCs and monocytes (Figure 2C).

**Figure 2:**
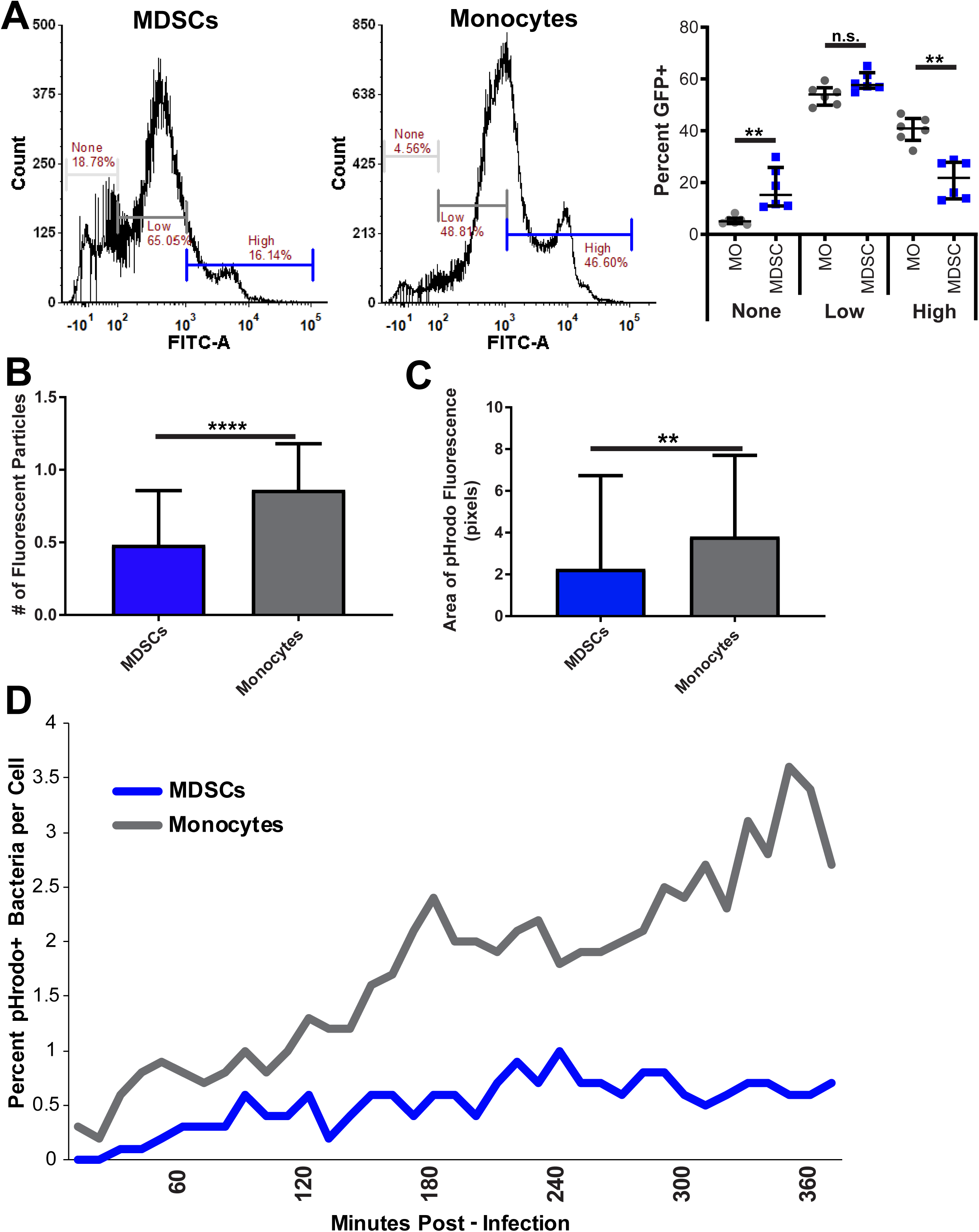
MDSCs are less efficient at bacterial uptake compared to monocytes. CD66^+^ MDSCs and CD14^+^ monocytes isolated from human umbilical cord blood PBMCs were infected with an MOI of 10 of Syto 9™- or pHrodo Red-labeled *E. coli* O1:K1:H7 and incubated at 37°C. For flow cytometry, cells were fixed in 4% paraformaldehyde and resuspended in PBS prior to collection. For imaging, cells were imaged and analyzed for pHrodo fluorescence using FIJI. For longitudinal imaging, cells were imaged every 10 minutes over a 6-hour period and quantified for pHrodo fluorescence. **(A)** Histogram and dot plots are representative of 2 independent experiments with three replicates per experiment. **(B and C)** Graphs are representative of 3 independent experiments. n = 63 and 70 images analyzed for MDSCs and monocytes, respectively. **(D)** The graph shown is representative of 2 independent experiments. n = 9 fields of view per cell type averaged at each time point per experiment. **(A)** Flow cytometry histograms and representative dot plot for MDSCs and monocytes display percent of cells that have not phagocytosed bacteria (none), have phagocytosed a low amount of bacteria (low), or have phagocytosed a high amount of bacteria (high). Grey circle symbols = monocytes, royal blue square symbols = MDSCs. **(B)** Quantification of the number of fluorescent bacterial particles phagocytosed by MDSCs and monocytes during infection. **(C)** Quantification of the area of pHrodo fluorescence in pixels phagocytosed by MDSCs and monocytes during infection. **(D)** Longitudinal phagocytosis of pHrodo bacteria during a 6-hour time course. Images were taken every 10 minutes of both MDSCs (blue line) and monocytes (grey line). Fluorescent bacteria per cell type were quantified at each time point from 9 fields of view. Statistical analyses in **(A, B, and C)** were performed using a Mann-Whitney U test. ** p ≤ 0.01, **** p ≤ 0.0001, n.s. not significant. Median with interquartile range displayed in all graphs.

To determine how quickly MDSCs and monocytes phagocytose bacteria, cells were infected and then longitudinally imaged over a 6-hour period. MDSC uptake of bacteria increased more gradually than that of monocytes and peaked at 4 hours (h) (Figure 2D). In contrast, internalization of bacteria by monocytes continued to increase more significantly through 6 h of infection. To determine whether MDSCs are able to eliminate bacteria at all or at a level comparable to monocytes, we implemented a previously described gentamicin protection assay (Gleave Parson et al., 2019). Briefly, MDSCs or monocytes were infected with bacteria for 1 h, and then treated with gentamicin to kill extracellular bacteria. Following 2 hours of gentamicin exposure, MDSCs and monocytes were permeabilized with 1% saponin at varying time points to quantify bacterial recovery. Previous work in our lab has shown that following 2 h of gentamicin treatment, nearly all bacteria are non-viable (Gleave Parson et al., 2019). We found a significantly higher bacterial recovery from MDSCs at 6- and 18- hours post gentamicin exposure (Figure 3). However, by 24 h, bacterial recovery from MDSCs was comparable to monocytes (Figure 3). Overall, these results suggest that although MDSCs are capable of bacterial uptake and elimination, they are significantly less efficient at these functions relative to monocytes.

**Figure 3:**
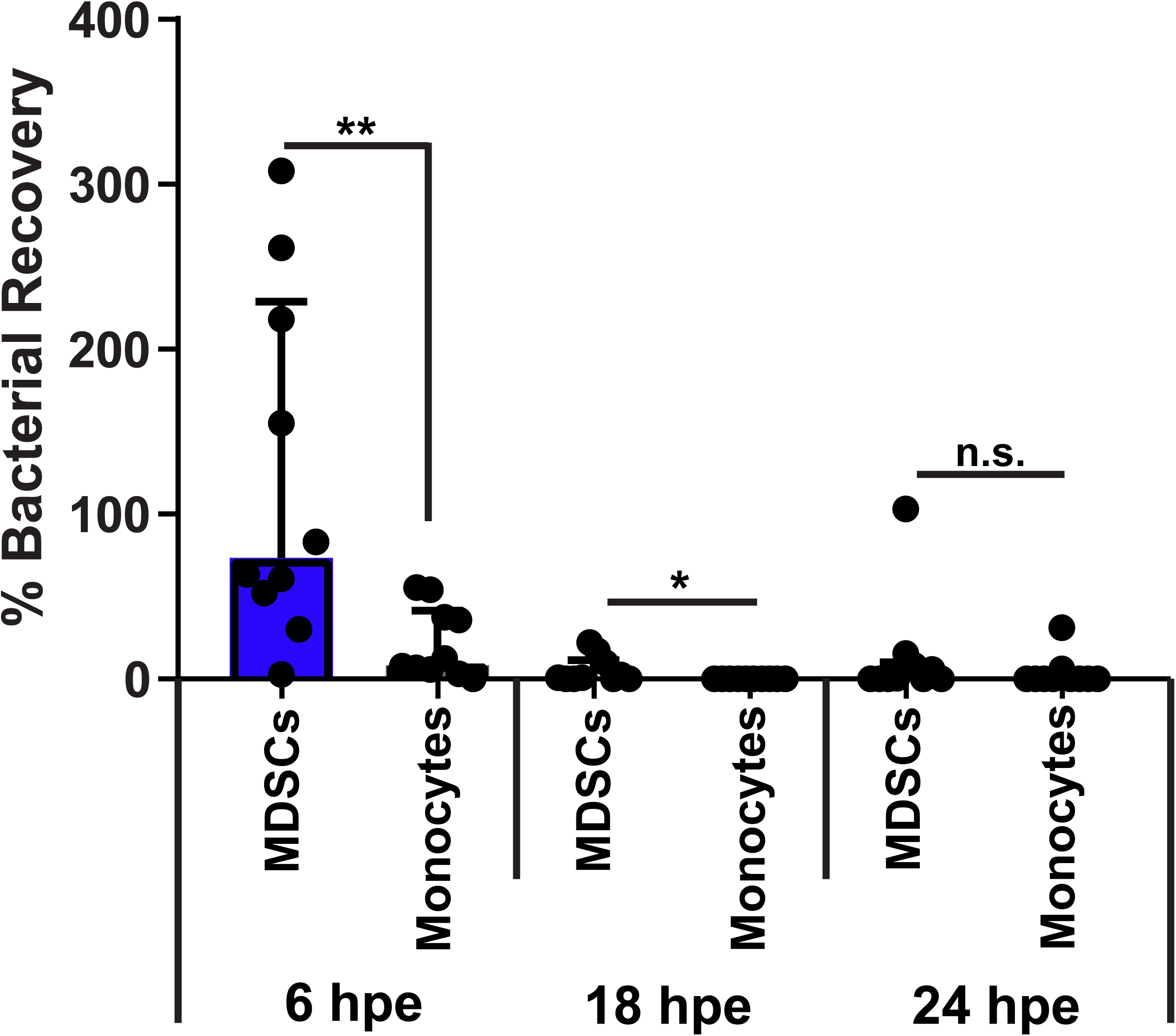
MDSCs are less efficient at bacterial elimination compared to monocytes. CD66^+^ MDSCs and CD14^+^ monocytes isolated from human umbilical cord blood PBMCs were infected with an MOI of 20 of *E. coli* O1:K1:H7 and incubated at 37°C for 1 hour. Media was replaced at this time point for gentamicin-supplemented media and cells were incubated for 2, 6, 18, and 24 hours post gentamicin exposure. At each time point, cells were permeabilized with 1% saponin, diluted ten-fold, and plated on TSA for standard bacterial enumeration. The graph represents bacterial recovery between MDSCs and monocytes at 6, 18, and 24 hours post exposure. All time points were normalized to the 2 hour time point. The data shown is representative of 5 independent experiments. Statistical analysis was performed using a Mann-Whitney U test. * p ≤ 0.05, ** p ≤ 0.01, n.s. not significant.

### Granulocytic MDSCs and monocytes express inflammatory cytokines during infection

Since MDSCs have some ability to eliminate pathogens during infection, we wanted to determine whether this is associated with a robust inflammatory response. To determine whether MDSCs express inflammatory cytokines at a level similar to monocytes during infection, we infected both cell types with *E. coli* and quantified TNFα expression at both the gene and protein levels. TNFα is a known biomarker for sepsis in patients and a classical proinflammatory cytokine produced by myeloid cells during infection (Samraj, Zingarelli, & Wong, 2013). We found that infected MDSCs are capable of expressing TNFα, but do not respond as robustly as infected monocytes (Figure 4A). Further, secreted cytokine levels were higher in infected monocytes compared to infected MDSCs. (Figure 4B). Overall, these data suggest that MDSCs express some level of inflammatory cytokines, but do not generate an inflammatory response comparable to monocytes during infection.

**Figure 4:**
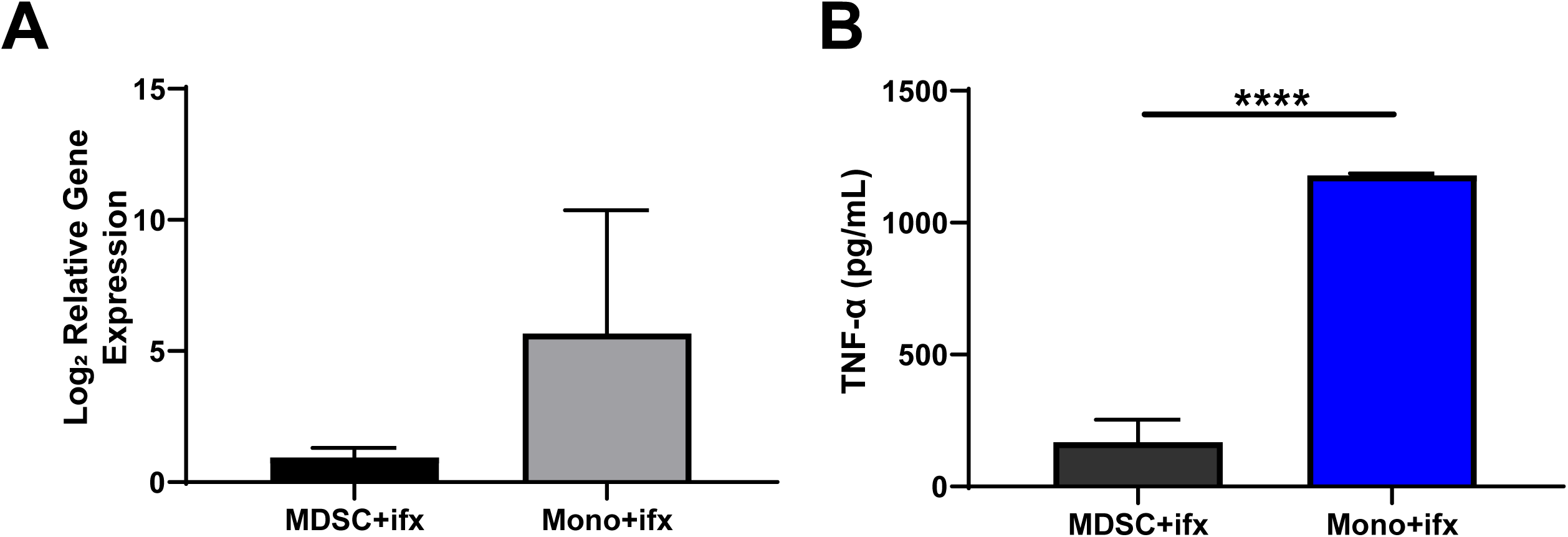
MDSCs and monocytes produce inflammatory cytokines during infection. CD66^+^ MDSCs and CD14^+^ monocytes isolated from human umbilical cord blood PBMCs were infected with an MOI 10 of *E. coli* O1:K1:H7 and incubated at 37°C for 6 hours. Supernatants were collected for inflammatory cytokine measurements. Cells were then lysed in TRI Reagent for RNA extraction, cDNA synthesis, and gene expression analysis of inflammatory cytokines. **(A)** Gene expression analysis of TNFα levels during infections. Infection levels were normalized relative to uninfected MDSC controls. Representative of 4 independent experiments, 10 replicates total per group. **(B)** Serum cytokine levels of TNFα measured via ELISA for infections of both MDSCs and monocytes. Cytokine levels were normalized based on standard curves for each protein. A representative of 2 independent experiments with 8 replicates total per group is shown. Statistical analysis was performed using an unpaired t-test for all panels. * p ≤ 0.05, **** p ≤ 0.0001.

### Granulocytic MDSCs release DNA during infection

To our surprise, time lapse imaging of MDSCs cultured with bacteria identified MDSC-generated thin extracellular structures that interacted with and connected to other MDSCs. Release of extracellular DNA traps is an important feature of granulocyte-mediated destruction of bacteria (Brinkmann et al., 2004; Papayannopoulos, 2018). Using the nucleic acid stain Sytox™ Green, we found high amounts of extracellular DNA strings released from what appear to be dead or dying MDSC cells since their cellular integrity is compromised as indicated by the availability of Sytox™ Green in the nuclear contents (Figure 5A). These strings were eliminated in the presence of DNAse I, demonstrating they are composed of nucleic acid (Figure 5A-B). Our quantification method demonstrated that there is a high amount of extracellular DNA in the absence of DNAse I (Figure 5B). However, because our method did not exclude DNA within dead or dying cells, the results did not achieve statistical significance between the untreated and DNAse I-treated groups (Figure 5B). To determine if MDSC extracellular (MeDNA) strings are important for bacterial elimination, we performed a gentamicin protection assay to enumerate intracellular killing along with standard plate counting of the culture supernatant to account for extracellular bacterial killing in the presence or absence of DNAse I. The extracellular and intracellular killing were combined to represent total levels of viable bacteria in Figure 6. We found that the addition of DNAse I to MDSC cultures did result in a trend toward increasing the total bacterial burdens (Figure 6). In contrast, total monocyte killing was not impacted by the addition of DNAse I (Figure 6). Similarly, when MDSCs were co-cultured with monocytes, the addition of DNAse I did not significantly alter the total bacterial killing (Figure 6). These results suggest that MeDNA may contribute to MDSC-dependent bacterial killing, but because of the relative inefficiency in elimination of bacteria relative to monocytes, in a mixed culture the contribution does not manifest as significant.

**Figure 5:**
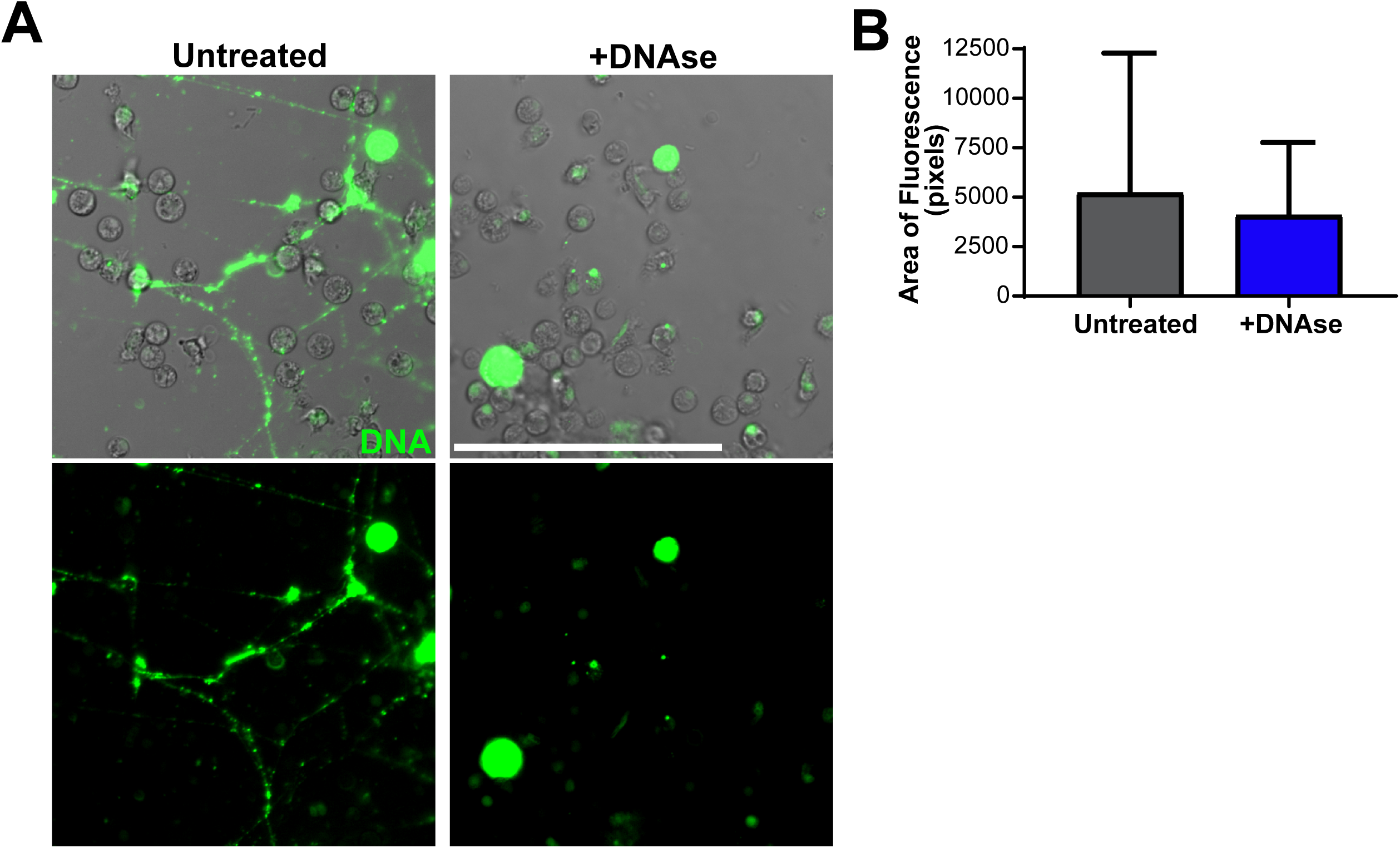
MDSCs produce extracellular DNA during infection. CD66^+^ MDSCs isolated from human umbilical cord blood PBMCs were infected with an MOI 10 of *E. coli* O1:K1:H7 and incubated at 37°C for ∼1.5 hours. To visualize extracellular DNA, 500 nM of Sytox™ Green was incorporated into cell media prior to imaging. To degrade DNA, 100 units of DNAse I was supplemented in media. Cells were imaged on a Nikon A1R confocal microscope at 20X and quantified for Sytox Green fluorescence in FIJI. **(A and B)** Representative images and quantification of Sytox Green fluorescence from 2 independent experiments are shown. n = 40 images per DNAse treated and untreated groups. In (B), black circle symbols = individual images for untreated cells, royal blue square symbols = individual images for DNAse-treated cells. Scale bar = 100 μm. Statistical analysis of **(B)** was performed using a Mann-Whitney U test; median with interquartile range is displayed.

**Figure 6:**
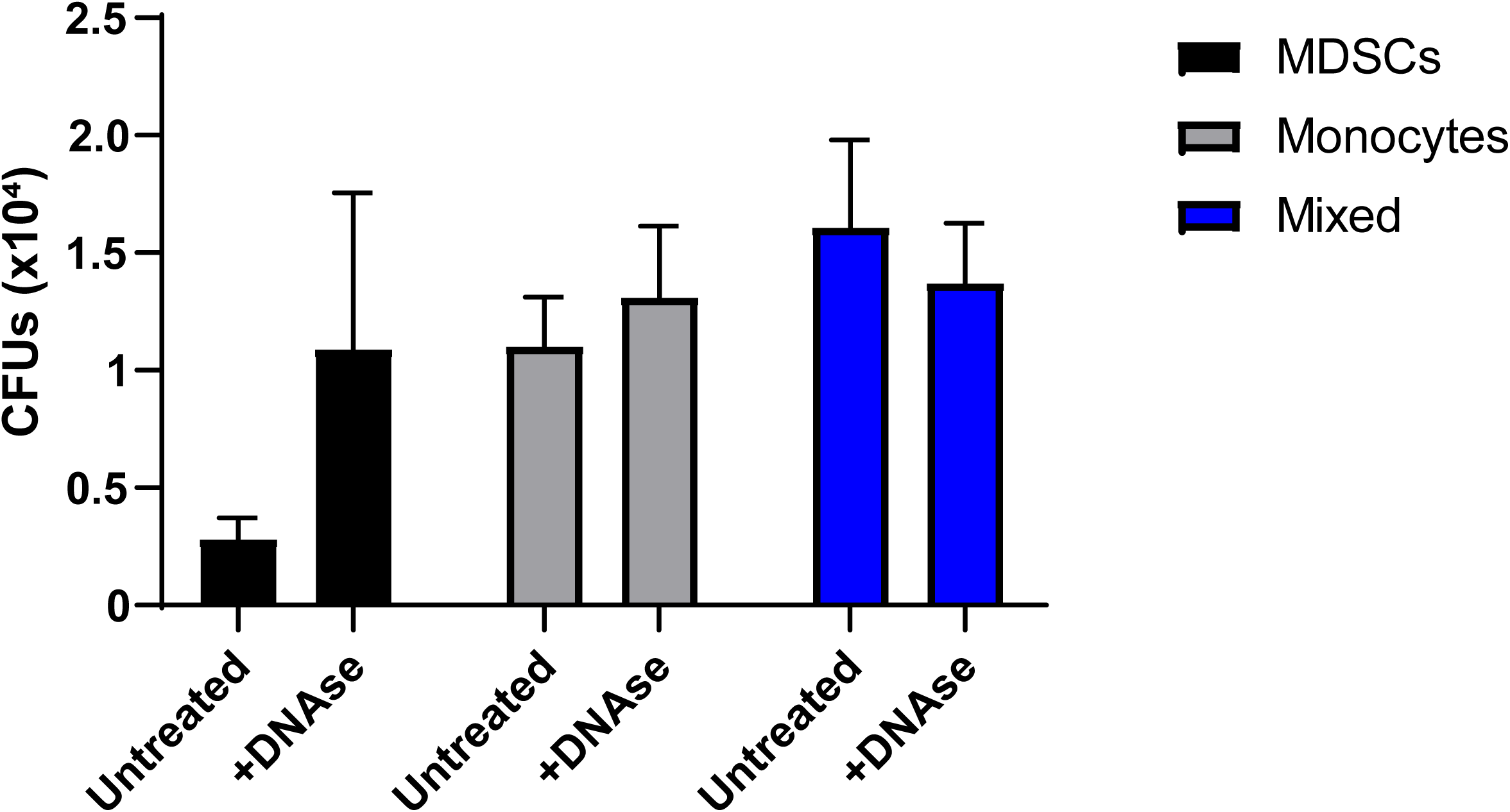
MDSC-derived extracellular DNA affects bacterial viability in MDSC-only cultures, but does not affect monocyte phagocytosis and killing of bacteria. CD66^+^ MDSCs and CD14^+^ monocytes isolated from human umbilical cord blood PBMCs were either single or co-culture infected with an MOI of 10 of *E. coli* O1:K1:H7 and incubated at 37°C for 6 hours. DNAse I (100 U) was added to appropriate cultures. For extracellular bacterial burdens, 50 μL of supernatant was collected from each infection, diluted ten-fold in PBS, and plated on TSA for standard plate counting. For intracellular burdens, 100 μL of 1% saponin was added to each well for 15 minutes, cell lysates were diluted ten-fold in PBS, and bacteria was plated on TSA for standard enumeration. Colony forming units (CFUs) of combined extracellular and intracellular bacteria in single or co-culture infections of MDSCs and monocytes untreated or treated with DNAse I at 6 hours post-infection. Statistical analysis was performed using a two-way ANOVA. *P*-values are as follows: interaction variation, p=0.3818, row factor variation (untreated vs. treated), p=0.4288, column factor variation (MDSCs vs. Monocytes vs. Mixed cultures), p=0.1156.

## Discussion

Myeloid-derived suppressor cells (MDSCs) have been well-studied in the context of cancer, but their direct involvement in host-pathogen interactions during infection has been less clear. Here, we describe one of the first in-depth studies on the direct interactions of MDSCs with bacteria. Our findings rigorously demonstrate that human granulocytic MDSCs have the ability to phagocytose and kill bacteria, although at a reduced efficiency compared to monocytes. In addition to these functions, we surprisingly observed release of DNA into the extracellular environment by MDSCs during infection. The extracellular DNA does promote MDSC-mediated bacterial killing, although the impact in a mixed cell culture is not yet clear. These activities are associated with a modest inflammatory response that does not rise to a level comparable with monocytes.

MDSCs phagocytose *E. coli* O1:K1:H7 in a dose-dependent manner. Using both flow cytometry and confocal microscopy, we were able to rigorously establish that MDSCs internalize bacteria and shuttle them into acidic compartments similar to professional phagocytes. We are the first to describe this pattern of intracellular trafficking in MDSCs. Leiber and colleagues previously reported that human granulocytic MDSCs could internalize bacteria (Leiber et al., 2017). Although their findings are in agreement with our results, they investigated the phagocytosis of a laboratory strain of *E. coli* at a single MOI of 50. Our study has evaluated bacterial internalization at a range of MOIs that includes low numbers of bacteria, and with a clinically relevant strain of *E. coli* responsible for invasive infections, such as sepsis and meningitis (Yao, Xie, & Kim, 2006). Additionally, two other studies have reported the internalization of *M. tuberculosis* and *M. bovis* BCG by MDSCs, again in agreement with our results (Agrawal et al., 2018; Magcwebeba et al., 2019; Martino et al., 2010). However, our study rigorously addressed the kinetics, frequency, and abundance of internalization using fluorescence microscopy. We further extended our approach to explore the fate of bacteria following internalization. He and colleagues reported enhanced killing of *E. coli* by human monocytic and granulocytic MDSCs from mice without procedural details that allow for complete interpretation (He et al., 2018). Based on these prior studies, we are the first to establish that bacteria internalized by MDSCs are trafficked to acidified compartments and further measure bacterial killing over time.

Our results demonstrate that MDSCs eliminate bacteria with reduced efficiency compared to monocytes. This analysis normalized the bacteria recovered to that which was internalized by each cell type at 2 hours post gentamicin treatment to account for differences in uptake, and allow for direct comparison of the rate of killing. Cellular mechanisms responsible for reduced internalization and killing by MDSCs compared to monocytes are currently unknown. Each function is independently impaired in MDSCs. For instance, since MDSCs are thought to be immature myeloid cells, the potential decrease in certain pattern recognition receptors (PRRs) on the cell surfaces of MDSCs could interfere in the ability of these cells to efficiently activate and internalize bacteria. One marker, CD14, is important in the recognition of lipopolysaccharide (LPS), a Gram-negative bacterial cell wall component (Devitt et al., 1998; Pugin et al., 1994). CD14 is highly expressed in monocytes and macrophages (Ost et al., 2016; Simmons, Tan, Tenen, Nicholson-Weller, & Seed, 1989), but the expression is low on granulocytic MDSCs (Supplemental Figure 1A). There are no reports of complement or mannose receptor expression by MDSCs that are frequently utilized for recognition and internalization of bacteria. These markers, as well as the presence or absence of other markers normally found in higher abundance on granulocytes and monocytes, could be partly responsible for the reduced efficiency in phagocytosis by MDSCs. To explain the reduced efficiency in bacterial clearance by MDSCs, it is possible that while bacteria are trafficked to lysosomes, this is done with reduced kinetics and efficiency. Acidified compartments within MDSCs may also have a more limited repertoire or abundance of hydrolytic molecules. Future studies will be necessary to address the cell biology of MDSCs and how it compares with professional phagocytes.

Granulocytic MDSCs release MeDNA during infection. These results are novel, as only one prior study shows that granulocytic MDSCs produce extracellular DNA (Alfaro et al., 2016). However, the gMDSCs described in that study do not inhibit T cell proliferation, a key characteristic of MDSCs, and thus may instead be low-density neutrophils (Rahman et al., 2019). At least some of the MeDNA appears microscopically to be associated with cell death; Sytox™ Green should not have access to the nucleus in cells that maintain membrane integrity. Lieber and colleagues (2017) reported an increased rate of apoptosis in MDSCs infected with *E. coli* (Leiber et al., 2017). Our observations are not consistent with an apoptotic form of cell death. The nature of this difference in findings is not clear, but may be influenced by the virulence of the bacteria. The MeDNA is associated with a reduction in bacterial viability in MDSC-only cultures, but the overall magnitude was not as striking as anticipated. Granule-packed neutrophils generate NETs that are strongly antibacterial (Brinkmann et al., 2004; P. Li et al., 2010). One possible explanation for limited bactericidal activity may be due to lower granule content in MDSCs compared to neutrophils (Rosales, 2018). Neutrophil granules are packed with highly antimicrobial contents, including defensins, cathepsins, and proteinases, and are released with extracellular DNA during NET formation (Borregaard, Sorensen, & Theilgaard-Monch, 2007). To our knowledge, there are no studies that describe the contents of the granules found in granulocytic MDSCs.

Experiments that explore the contents and abundance of these granules found in granulocytic MDSCs compared to neutrophils will help us improve our understanding of the limited toxicity of MeDNA. Extracellular DNA from neutrophils has been implicated in endothelial cell damage and death (Clark et al., 2007; Gupta et al., 2010). Additionally, extracellular histones, like the citrullinated histone H3, cause damage to multiple cell types and can lead to organ failure (Kutcher et al., 2012). Histone release during sepsis promotes endothelial cell dysfunction, hypoxia in tissues, and cell death (Wildhagen et al., 2014; Xu et al., 2009). Since the co-culture of MDSCs with monocytes did not significantly alter clearance of bacteria, this finding suggests that MeDNA does not compromise monocyte function or viability. Additional studies will be needed to further evaluate if MeDNA is cytotoxic to other cell types.

In conclusion, our body of work demonstrates evidence of MDSC phagocytic capacity during acute infection. This is not associated with a robust inflammatory response as compared with monocytes. Additionally, we have unexpectedly discovered that MDSCs release extracellular DNA that has modest antibacterial activity in a homogenous culture of MDSCs, but is less influential during co-culture with monocytes. Ongoing work in our lab continues to characterize these mechanisms to further understand how MDSCs directly interact with bacteria and other immune cells during infection. These newly discovered MDSC activities may direct the future use of novel therapies to improve neonatal immunity and disease outcome during severe infections.

## Materials and Methods

### Cell Culture

Human umbilical cord blood was obtained from the Cleveland Cord Blood Center under West Virginia University Institutional Review Board (IRB) approval. Blood was donated from healthy infants of gestational age ≥ 37 weeks. All donors are anonymous and de-identified. Whole blood was centrifuged at 1500 x g for 10 minutes to obtain buffy coats. These were further subjected to Ficoll (GE Healthcare Life Sciences, Chicago, IL) density gradient centrifugation at 400 x g for 30 minutes to isolate peripheral blood mononuclear cells (PBMCs). CD66abce^+^ MDSCs, CD14^+^ monocytes, and CD4^+^ T cells were isolated by immunomagnetic selection using their respective Miltenyi Biotec isolation reagents (Miltenyi Biotech, Bergisch Gladbach, Germany). MDSCs and monocytes were incubated at a concentration of 1-7×10^5^ cells/well in FluoroBrite Dulbecco’s Modified Eagle Medium (DMEM, ThermoFisher Scientific, Waltham, MA) supplemented with 10% human serum, 25 mM HEPES, and 2 mM L-glutamine. T cells were cultured at a concentration of 1×10^5^ cells/well in RPMI-1640 (Mediatech, Manassas, VA), supplemented with 10% human serum, 100 U/mL penicillin/streptomycin, 2 mM L-glutamine, 25 mM HEPES, 1 mM sodium pyruvate, and 0.05 mM 2-mercaptoethanol. Human cultures were incubated at 37°C in 48- or 96-well plastic bottom plates, or in a 35 mm ibidi Quad μ-Dish (ibidi, Fitchburg, WI) for confocal/epifluorescence imaging.

### Fluorescent Labeling and Bacterial Infection

Human MDSCs and monocytes were infected with an MOI ∼2-200 of *Escherichia coli* strain O1:K1:H7. The bacteria were taken from pre-titered frozen cultures and washed 1x in phosphate-buffered saline (PBS; Corning, Manassas, VA), centrifuged at 12,000 x rpm, and resuspended in a volume equivalent to an inoculum of 50 μL/well. Bacteria were labeled with 5 μM Syto™ 9 Green Fluorescent Nucleic Acid Stain or 500 nM pHrodo Red SE (ThermoFisher Scientific, Waltham, MA) and washed 3-5 times with PBS prior to infection. For extracellular bacterial recovery assays, bacteria were taken directly from culture supernatants, diluted ten-fold in PBS, and enumerated by standard plate counting on tryptic soy agar (TSA; Becton, Dickinson and Company, Sparks, MD) incubated at 37°C overnight. To visualize extracellular DNA, 500 nM of Sytox™ Green Nucleic Acid Stain (ThermoFisher Scientific, Waltham, MA) was added prior to microscopy.

### Gentamicin Protection Assay

Human MDSCs and monocytes were infected at an MOI of 20 for 1 hour at 37°C. At 1 hour post-infection, supernatants were discarded and cells were supplemented with new media containing 100 μg/mL gentamicin (Quality Biological, Gaithersburg, MD) to eliminate extracellular bacteria. Cells were incubated for 2, 6, 18, and 24 hours and then permeabilized using 1% saponin in PBS (MP Biomedicals, Solon, OH). Cell lysates were diluted ten-fold in PBS and bacteria was enumerated by standard plate counting. In experiments incorporating DNAse I, media was supplemented with 100 units of DNAse I (Roche, Basel, Switzerland) during initial incubation and media replacement with gentamicin.

### Flow Cytometry

Cells and bacteria were incubated at 37°C for varying time points. At each time point, cells were collected in 500 μL of 4% paraformaldehyde (Affymetrix, Cleveland, OH) and kept at 4°C until use. Cells were resuspended in 400 μL PBS and approximately 10,000 events were collected on an LSRFortessa (Becton, Dickinson and Company, Sparks, MD). Percent cells gated in FITC- (488 laser, 490/525 nm excitation/emission), PE- (561 laser, 496/578 nm excitation/emission), or Pacific Blue-(405 laser, 410/455 nm excitation/emission) channels were used for data analysis. For cell marker profiling, MDSCs were immunolabeled with PE-conjugated anti-HLA-DR (Invitrogen, San Diego, CA), anti-human CD66b-PE (BioLegend, San Diego, CA), FITC-conjugated anti-CD14 (R&D Systems, Minneapolis, MN), and FITC-conjugated anti-CD33 (BioLegend, San Diego, CA).

### T Cell Proliferation Assay

CD4^+^ T cells were labeled with 5 μM CellTrace Violet (ThermoFisher Scientific, Waltham, MA) for 20 minutes at 37°C. The labeling was quenched with 10% fetal bovine serum (FBS; Gemini Bio-Products, West Sacramento, CA) in PBS and cells were resuspended in T cell media. Cells were then plated at 1×10^5^ cells/well in a non-tissue culture treated 96-well plate (Corning, Corning, NY) and incubated for 2 hours at 37°C. CD66^+^ MDSCs were then added at a concentration of 1×10^5^ cells/well. To promote proliferation, the culture was supplemented with 100 units of interleukin-2 (IL-2; Shenandoah Biotech, Warwick, PA) or 3×10^5^ CD3/CD28 Dynabeads/well (ThermoFisher Scientific, Waltham, MA). Cells were incubated for up to 4 days at 37°C. Cells were harvested on days 1, 2, 3, and 4 and fixed in 4% paraformaldehyde until analysis via flow cytometry.

### Quantitative Real Time PCR

MDSCs and monocytes were cultured with or without *E. coli* at an MOI ∼10 in a 48-well plastic-bottom plates for 6 hours. At 6 hours, cells were resuspended in 200 μL of TRI Reagent (Molecular Research Center, Cincinnati, OH). Following phase separation, the aqueous layer was mixed with an equal volume of cold 75% ethanol and transferred to MicroElute LE RNA Columns (Omega Bio-Tek, Norcross, GA). After RNA extraction, first-strand cDNA was synthesized using iScript reagents (Bio-Rad, Hercules, CA). Quantitative PCR reactions included cDNA diluted 2-fold from synthesis, gene-specific Taqman primer probe sets (Applied Biosystems, Foster City, CA), and iQ Supermix (Bio-Rad, Hercules, CA). Cycling was performed in duplicate using a Step One Plus™ Real Time detection system (Applied Biosystems, Foster City, CA). Gene-specific amplification was normalized to GAPDH as an internal reference gene and gene expression was normalized relative to uninfected MDSC controls using the formula 2^-ΔΔCt^.

### Cytokine Measurements

Supernatants from infections were collected and clarified by standard techniques. TNFα protein levels were measured in duplicate or triplicate using a Ready-Set-Go! ELISA kit (eBioscience, San Diego, CA). Results were analyzed and normalized to standard curves for the cytokine and concentrations of cytokine was graphed accordingly.

### Fluorescence Microscopy

A Nikon A1R confocal microscope was used for confocal and epifluorescence imaging (Nikon, Melville, NY). Objective lenses with powers of 20X (numerical aperture [NA], 0.75), 40X (oil, NA, 1.3), and 60X (oil, NA, 1.4) were used. Images are overlays of differential interference contrast (DIC) and fluorescence images or fluorescence image panels. Syto 9™/Sytox Green™ and pHrodo Red were detected by optical lasers/filters for excitation/emission at 490/525 nm (FITC) and 555/580 nm (TRITC), respectively. Images were analyzed in ImageJ (FIJI, www.fiji.sc). Briefly, images were thresholded for bacterial fluorescence, and each area was quantified with identical settings per experiment and processed identically. For 6-hour time lapses, cells were imaged on a Lionheart FX automated microscope (BioTek, Winooski, VT). Images were acquired using a 20X objective (NA, 0.45) and analyzed using Gen5 Image+ software (version 3.05.11; BioTek, Winooski, VT).

### Statistical Analysis

All statistical analyses were performed using GraphPad Prism software (version 8; La Jolla, CA). Data was tested using non-parametric or parametric measures, as indicated in the figure legends.

## Acknowledgements

We would like to thank members of the Robinson lab for stimulating discussions, Dr. Matthew Daddysman for help with Gen5 Image+ image analysis, Drs. Amanda Ammer and Karen Martin for microscopy guidance, and Dr. Kathy Brundage for flow cytometry guidance. This work was supported by West Virginia University Institutional funds.

## Figure Legends

**Supplemental Figure 1:**
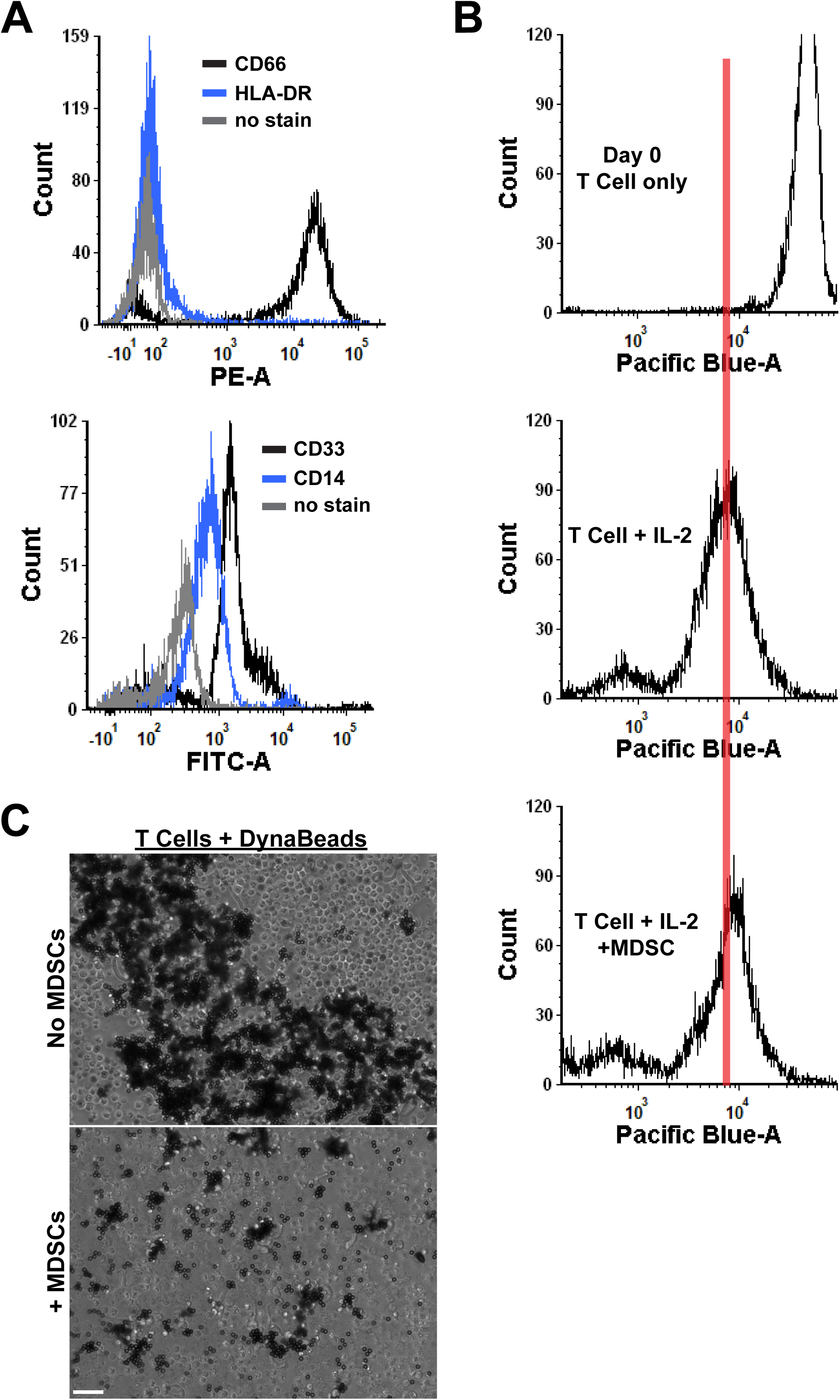
Neonatal human MDSCs have characteristic cell surface markers and suppress T cell proliferation. CD66^+^ MDSCs from human umbilical cord blood PBMCs were either labeled with cell surface markers for cell marker profiling or were co-cultured at a 1:1 ratio with CD4^+^ T cells for 4 days for a T cell proliferation assay. For cell marker profiling, MDSCs were labeled with antibodies for CD66, HLA-DR, CD33, and CD14, fixed in 4% paraformaldehyde, and resuspended in PBS prior to collection on the flow cytometer. For T cell proliferation assays, MDSCs were incubated with T cells stimulated with IL-2 (100 U) for 4 days. Cells were collected each day and fixed in 4% paraformaldehyde for flow cytometry analysis. Cells supplemented with CD3/CD28 DynaBeads were imaged on a Lionheart FX automated microscope to visualize clonal expansion of T cells surrounding beads during proliferation. **(A)** Shown are representative histogram overlay plots of cell surface markers for MDSCs compared to no stain controls. The top panel shows PE-labeled CD66 and HLA-DR expression on cell surfaces. The bottom panel shows FITC-labeled CD33 and CD14 expression on cell surfaces. Shifts to the right represent increasing fluorescence. Black lines = CD66 or CD33 expression in top and bottom panels, respectively, blue lines = HLA-DR or CD14 expression in top and bottom panels, respectively, grey lines = no stain control in both panels. A representative histogram of 2 independent experiments is shown. **(B)** Shown are representative histogram plots of T cell proliferation. Stimulated T cells at Day 0, IL-2 stimulated T cells at day 3, and IL-2 stimulated T cells supplemented with MDSCs at day 3 are displayed. The red vertical line on all plots is used to visualize the shift in proliferation in all plots. The data shown is representative of 5 independent experiments. **(C)** Representative DIC images of T cells supplemented with DynaBeads ± MDSCs are shown. Black coverage is representative of beads associated with proliferating T cells. The data shown is representative of 5 independent experiments. Scale bar = 100 μm.

